# Far-red fluorescent genetically encoded calcium ion indicators

**DOI:** 10.1101/2020.11.12.380089

**Authors:** Rochelin Dalangin, Mikhail Drobizhev, Rosana S. Molina, Abhi Aggarwal, Ronak Patel, Ahmed S. Abdelfattah, Yufeng Zhao, Jiahui Wu, Kaspar Podgorski, Eric R. Schreiter, Thomas E. Hughes, Robert E. Campbell, Yi Shen

## Abstract

Genetically encoded calcium ion (Ca^2+^) indicators (GECIs) are widely-used molecular tools for functional imaging of Ca^2+^ dynamics and neuronal activities on a single cell level. Here we report the design and development of two new far-red fluorescent GECIs, FR-GECO1a and FR-GECO1c, based on the monomeric far-red fluorescent protein mKelly. We characterized these far-red GECIs as purified proteins and assessed their performance when expressed in cultured neurons. FR-GECOs have excitation and emission maxima at ~ 596 nm and ~ 644 nm, respectively, display large responses to Ca^2+^ (Δ*F*/*F*_0_ = 6 for FR-GECO1a, 18 for FR-GECO1c), and are bright under both one-photon and two-photon illumination. FR-GECOs also have high affinities (apparent *K*_d_ = 29 nM for FR-GECO1a, 83 nM for FR-GECO1c) for Ca^2+^, and they enable sensitive and fast detection of single action potentials in neurons.

## Introduction

A property of most mammalian tissues is that they are most transparent to wavelengths of light between ~600 nm and ~1300 nm (the range often referred to as the “optical window”)^1,2^. This wavelength range falls between the absorbance profile of hemoglobin, which predominates at wavelengths below 600 nm, and the absorbance profile of water, which predominates at wavelengths greater than 1300 nm. Due to the greater tissue transparency in this wavelength range, fluorescent probes that absorb and emit efficiently within the optical window are highly desirable for *in vivo* imaging. In addition, fluorescent probes with longer excitation wavelengths are associated with lower phototoxicity and autofluorescence, reduced crosstalk with green fluorescent indicators, and better spectral compatibility with blue or cyan light-activated optogenetics tools.

Realization of the advantages of fluorophores that excite and emit within the optical window has been a driving force for molecular tool engineers to shift the excitation and emission wavelengths of genetically encodable fluorophores, such as standard red fluorescent proteins (RFP) with excitation maxima (λ_ex_) at 550 to 580 nm and emission (λ_em_) at 580 to 620 nm, into the far-red region of the spectrum. This longstanding effort has yielded a plethora of far-red FPs (**Figure 1A**) with λ_ex_ > 580 nm and λ_em_ > 620 nm (Refs. 3–12). Efforts to engineer biliverdin (BV)-binding FPs, that fluoresce at even further red-shifted wavelengths, have resulted in near infrared (NIR) FPs with λ_ex_ > 640 nm and λ_em_ > 670 nm (Refs. 13–15). The key difference between these two classes of genetically encodable fluorophores is that the red and far-red FPs autocatalytically form their own chromophore and are homologous with GFP, but genetically encoded NIR FPs use the biliverdin cofactor as chromophore.

**Figure 1.**
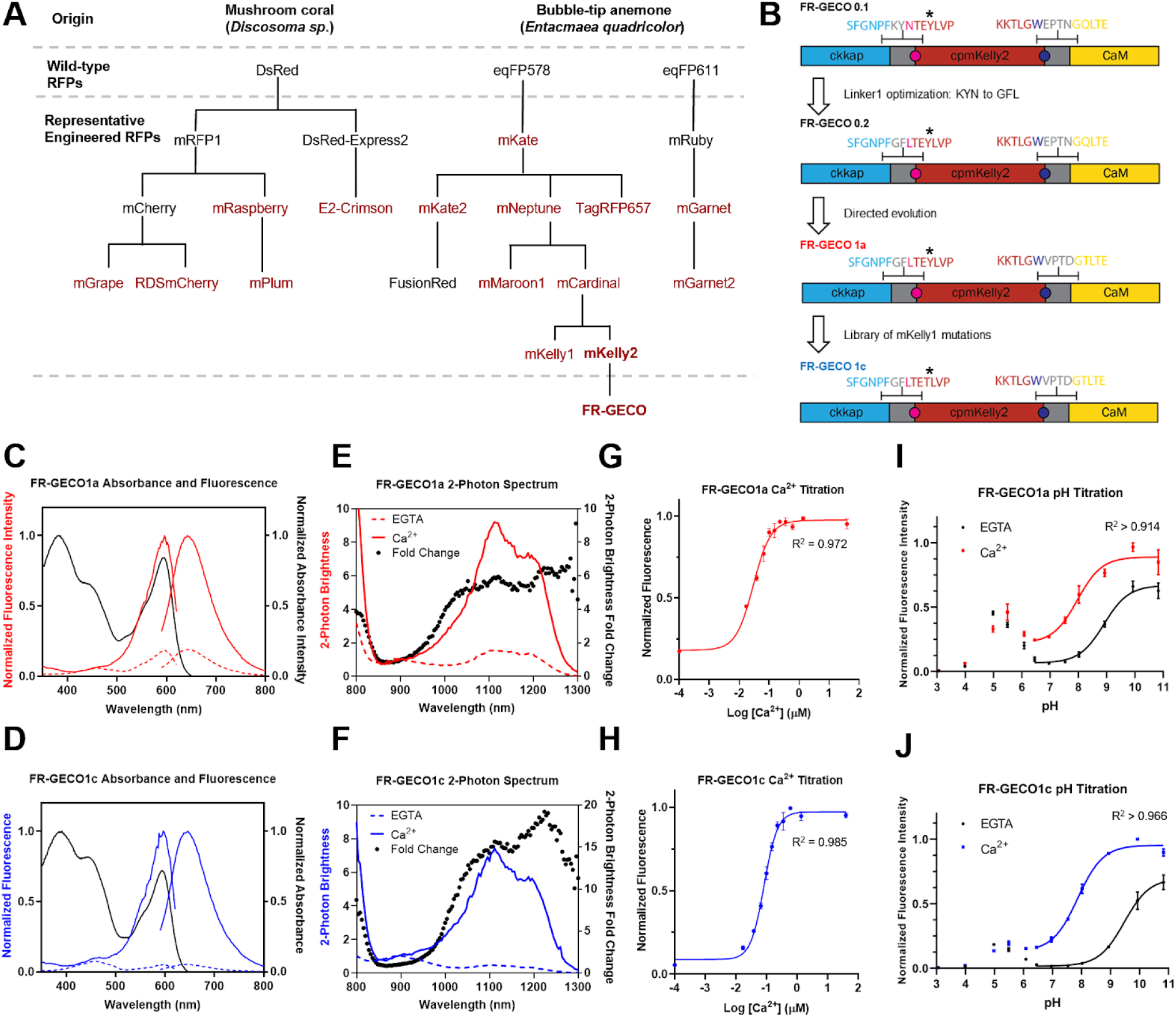
Engineering and characterization of FR-GECO1a and FR-GECO1c. **A**. Selected far-red FPs and FR-GECO genealogy. Far-red FPs (λ_em_ > 620 nm) are highlighted in red. **B**. Schematic illustration of FR-GECO1 design and engineering. Linker residues are in grey; asterisk indicates position 35. Magenta and blue colored amino acids (and correspondingly colored circles in the schematic), represent the positions of ‘gatepost’ residues that define the first and last residues of cpmKelly2. Absorbance, excitation (dashed) and emission (solid) spectra of FR-GECO1a (**C**) and FR-GECO1c (**D**). Two-photon spectra and dynamic range of FR-GECO1a (**E**) and FR-GECO1c (**F**). *K*_d_ titration (n = 3) with FR-GECO1a (**G**) and FR-GECO1c (**H**). pH sensitivity (n = 3) of FR-GECO1a (**I**) and FR-GECO1c (**J**). Error bars represent S.E.M.

Genetically encodable fluorophores can be engineered into genetically encoded indicators. One of the most important examples is the genetically encoded Ca^2+^ indicator (GECI) which can be used for detection and imaging of cell signalling and neuronal activities. The latest generation of red GECIs for neuronal activity detection include jRCaMP1a/b, jRGECO1a, K-GECO1, and XCaMP-R, all with single-photon excitation and emission peaks outside of the optical window^16–18^). Among currently available RFP-based GECIs, the most red-shifted variant that uses a fluorescent protein with an autocatalytic chromophore is CAR-GECO1 (λ_ex_ ~ 560 nm, λ_em_ ~ 609 nm)^19^). Near-infrared GECI indicator NIR-GECO1 (λ_ex_ ~ 678 nm, λ em ~ 704 nm) based on the BV-binding FP mIFP has been recently reported. Although the spectrum of NIR-GECO1 lies well within the optical window, the molecular brightness of NIR-GECO1 is relatively dim^20^).

In an effort to achieve the ideal compromise between red-shifts in the excitation and emission maxima and brightness, we have undertaken the development of two bright intensiometric far-red fluorescent GECI variants, FR-GECO1a (λ_ex_ ~ 596 nm, λ_em_ ~ 642 nm) and FR-GECO1c (λ_ex_ ~ 596 nm, λ_em_ ~ 646 nm), based on the recently engineered far-red FP mKelly1 (λ_ex_ ~ 596 nm, λ_em_ ~ 656 nm) and mKelly2 (λ_ex_ ~ 598 nm, λ_em_ ~ 649 nm)^12^). These genetically encoded far-red Ca^2+^ indicators will open new avenues for multicolor Ca^2+^ imaging in combination with other optogenetic indicators and actuators, as well as functional Ca^2+^ imaging in deep tissue *in vivo*.

## Results

### Protein engineering

Initial efforts to engineer a far-red Ca^2+^ indicator followed two parallel strategies. The first strategy was to graft key mutations for far-red fluorescence onto existing red GECIs. We used R-GECO1 (Ref. 21), CH-GECO1 (Ref. 22) and K-GECO1 (Ref. 17) as templates and introduced key mutations from E2-Crimson (λ_ex_ ~ 611 nm, λ_em_ ~ 646 nm, tetrameric)^5^), RDSmCherry (λ_ex_ ~ 600 nm, λ_em_ ~ 630 nm)^7^), and mNeptune (λ_ex_ ~ 600 nm, λ_em_ ~ 650 nm)^4^), respectively (**Figure 1A, Supplementary Table 1**). Unfortunately, complete loss of fluorescence or no substantial spectral red-shifts were observed in these designed prototypes. The second strategy was *de novo* engineering of a GECI starting from far-red FP scaffolds. However, our attempts to engineer GECI based on mNeptune^4^) and mCardinal^8^) did not yield fluorescent prototypes. As both mNeptune and mCardinal retained weak dimerization tendencies^4,8,12,23^, we suspected that the failure of these prototypes to fluoresce could be due to the circular permutation site and/or insertion site of calmodulin and its binding peptide overlapping with the oligomerization interface, possibly disrupting dimerization that was crucial to the proper folding and/or chromophore maturation of these proteins.

mKelly1 and mKelly2 are far-red FP variants of mCardinal that were engineered to have increased monomericity by a combination of deletion of the C-terminal “tail”, directed evolution, and consensus design^12^) (**Figure 1A**). We rationalized that the enhanced monomericity of mKelly1 and mKelly2 might facilitate circular permutation and further engineering to create GECIs. We thus circularly permuted (cp) mKelly2, the brighter of the two mKelly variants, at Thr143 (numbering according to the crystal structure for mCardinal^8^); PDB ID: 4OQW). This designed cpmKelly2 was then used to replace the cpFusionRed RFP in K-GECO1 (Ref. 17). K-GECO1 was chosen as the starting scaffold because the use of the ckkap peptide instead of RS20 as the calmodulin binding peptide yielded a sensor with high sensitivity, strong affinity, good linearity and fast kinetics. Additionally, it is currently the only GECI engineered using a FP variant derived from eqFP578, from which mKelly2 was also derived (**Figure 1A**). This initial construct resulted in a dimly fluorescent variant we named FR-GECO0.1 (**Figure 1B**). Optimization of linker1 (the amino acids linking the ckkap peptide to cpmKelly2) led to the identification of an improved, but still dim, variant, FR-GECO0.2, with λ_ex_ = 586 nm, λ_em_ = 632 nm and ~7-fold increase of fluorescence upon Ca ^2+^ addition. We then subjected FR-GECO0.2 to eight rounds of iterative directed evolution to improve its brightness and Ca^2+^ response while ensuring that all the variants chosen to be used as gene templates for the next round maintained a far-red emission peak (**Figure 1B**). Ultimately, our efforts led to a bright and sensitive far-red GECI, FR-GECO1a, which incorporated 19 mutations relative to the initial FR-GECO0.1 gene (**Supplementary Table 2**). FR-GECO1a had λ_ex_ = 596 nm, λ_em_ = 642 nm, and exhibited a 6-fold increase in fluorescence intensity in the presence of Ca^2+^(**Figure 1C**).

With the mKelly2-derived FR-GECO1a in hand, we attempted to incorporate mKelly1-specific mutations in an effort to achieve a slightly more red-shifted emission peak and higher photostability^12^). Notably, mKelly1 was engineered from mCardinal in parallel with mKelly2, and differed from mKelly2 by six mutations. To test whether these mutations could further improve FR-GECO1a, we screened a combinatorial library of FR-GECO1a containing all possible combinations of the six mKelly1-specific mutations. From this library we isolated a variant with λ_ex_ = 596 nm and λ_em_ = 646 nm. It exhibits an 18-fold increase in fluorescence intensity in the Ca^2+^-bound state, which is an over 2-fold improvement relative to that of FR-GECO1a. This variant, which we termed FR-GECO1c (c for **c**ontrast) incorporated a single mutation, Tyr35Thr (numbering according to FR-GECO1a sequence, **Figure 1B** and **Supplementary Figure 1**) which is positioned in close proximity to the first gatepost residue Leu32 (**Figure 1B**).

### *In vitro* characterization

To characterize the photophysical properties of FR-GECO1a and FR-GECO1c, we purified the bacterially-expressed protein. Spectral analysis of the purified proteins revealed that the 596 nm excitation peak of both FR-GECO1a and FR-GECO1c were slightly blue-shifted by 2 nm relative to mKelly2 (598 nm), and that the emission peaks of the FR-GECO1a (642 nm) and FR-GECO1c (644 nm) were also slightly blue-shifted relative to mKelly2 (649 nm) (**Figure 1C, D** and **Table 1**). The one-photon (1P) molecular brightness of FR-GECO1a (9.16) and FR-GECO1c (9.35) in their Ca^2+^-bound states are approximately 20% brighter than their template FP, mKelly2 (7.7) (**Table 1**). These results were attributed to a near doubling of the quantum yield (QY, Φ), which offset the effect of the indicators’ smaller effective extinction coefficients (EC, ε) compared to mKelly2. In the Ca^2+^-free and Ca^2+^-bound states, FR-GECO1a has ECs of 5,710 and 27,000 M^−1^cm^−1^, respectively, and FR-GECO1c has ECs of 4,850 and 26,500 M^−1^cm^−1^. The two-fold increase in the Ca^2+^-dependent Δ*F*/*F*_0_ of FR-GECO1c relative to FR-GECO1a (attributable to the Tyr35Thr mutation), was found to result primarily from the 2-fold smaller QY in the Ca^2+^-free state of FR-GECO1c.

**Table 1.** Photophysical characterization of FR-GECO1a, FR-GECO1c, and mKelly2.

| Protein Name | FR-GECO1a |  | FR-GECO1c |  | mKelly2* |
| --- | --- | --- | --- | --- | --- |
|  | Ca<sup>2+</sup>-free | Ca<sup>2+</sup>-bound | Ca<sup>2+</sup>-free | Ca<sup>2+</sup>-bound |  |
| Excitation maxima $\lambda_{ex}$ (nm) | 596 | | 596 | | 598 |
| Emission maxima $\lambda_{em}$ (nm) | 642 | | 646 | | 649 |
| Ca<sup>2+</sup>-dependent $\Delta F/F_0$ | $5.8 \pm 0.2$ | | $18.1 \pm 0.4$ | | n/a |
| $K_d$ (nM) | 29 | | 83 | | n/a |
| Apparent Hill coefficient ($n_H$) | 1.49 | | 1.98 | | n/a |
| Apparent $pK_a$ | 8.92 | 7.96 | 9.47 | 7.88 | 5.6 |
| $k_{on}$ (M<sup>-1</sup>s<sup>-1</sup>) | $9.07 \times 10^4$ | | $5.31 \times 10^6$ | | n/a |
| Rise $t_{1/2}$ (μs) | 7.64 | | $1.31 \times 10^{-2}$ | | n/a |
| $k_{off}$ (s<sup>-1</sup>) | 3.711 | | 6.393 | | n/a |
| Decay $t_{1/2}$ (ms) | 269 | | 156 | | n/a |
| Extinction Coefficient $\epsilon$ (M<sup>-1</sup>cm<sup>-1</sup>) | 5,710 | 27,000 | 4,850 | 26,500 | 43,000 |
| Quantum Yield $\Phi$ | 0.19 | 0.34 | 0.077 | 0.35 | 0.18 |
| Brightness ($\times 1000$) | 1.08 | 9.16 | 0.375 | 9.35 | 7.7 |
| 2P Brightness $F_2$ (GM),<br>$\lambda_{max}$ in parentheses | 1.56<br>(1112 nm) | 9.2<br>(1112 nm) | 0.48<br>(1108 nm) | 7.4<br>(1112 nm) | n/d |
| $\Delta F/F_0$ (2P), $\lambda_{max}$ in parentheses | 5.9 (1112 nm) | | 15.4 (1112 nm) | | n/a |

\*Values from Wannier *et al.*<sup>12</sup>. Errors represent S.E.M. n/a: not applicable. n/d: not determined.

Under two-photon (2P) conditions, both FR-GECO1a and FR-GECO1c had peak 2P excitation at 1112 nm (**Figure 1E, F**), with FR-GECO1a having a ~ 25% higher 2P molecular brightness (9.2 GM) than FR-GECO1c (7.4 GM) in the Ca^2+^-bound state (**Table 1**). Both FR-GECO1a and FR-GECO1c, in Ca^2+^-saturated states *in vitro*, were among the brightest red GECIs upon 2P excitation^24^) (**Supplementary Table 3**). The optimal wavelength range for 2P excitation, with close to maximum fluorescence change, was between 1050 and 1200 nm (**Figure 1E, F**). The overall Δ*F*/*F*_0_ of FR-GECO1a and FR-GECO1c under 1P and 2P conditions were comparable. Taken together, FR-GECO1a and FR-GECO1c exhibited bright far-red fluorescence under both 1P and 2P illumination.

Consistent with other ckkap-based GECIs^17,18,25^, static Ca^2+^ titrations revealed that both FR-GECO1a and FR-GECO1c have relatively high affinities for Ca^2+^ and Hill coefficients (n_H_) greater than unity (**Figure 1G, H**). FR-GECO1a (*K*_d_ = 32 nM; n_H_ = 1.49) has a substantially lower *K*_d_ and n_H_ than FR-GECO1c (*K*_d_ = 87 nM; n_H_ = 1.98). These relatively high Ca^2+^ binding affinities suggested FR-GECO1a and FR-GECO1c are well-suited for neuronal activity detection. Kinetic measurements revealed fast fluorescence responses from Ca^2+^ association and dissociation for both FR-GECO1a and FR-GECO1c (**Table 1, Supplementary Figure 2**). Specifically, FR-GECO1a (half decay time = 269 ms) and FR-GECO1c (half decay time = 156 ms) are among the fastest decaying GECIs with high Ca^2+^ affinity^26^). The pH sensitivity measurements (**Figure 1I, J**) indicated that both FR-GECO1 variants undergo a shift in p*K*_a_ upon binding Ca^2+^with a p*K*_a_ of ~ 9 in the Ca^2+^-free state and ~ 8 in the Ca^2+^-bound state. Both variants exhibited a small peak at pH = 5 in the Ca^2+^-free state and 5.5 in the bound state, suggesting multiple ionizable groups, which resembles the pH characterization of CH-GECO series of red GECIs^27^).

To test whether FR-GECOs can be photoactivated by blue light (as observed for the R-GECO family of red GECIs and, to a lesser extent, K-GECO1)^16,17,19^, purified proteins were imaged using 561 nm (3.70 W/cm^2^)) excitation laser with 488 nm (0.71 W/cm^2^)) blue light laser pulses (50 ms). FR-GECO1a and FR-GECO1c were found to exhibit moderate increases in fluorescence emission with blue light illumination, in both the Ca^2+^-bound and Ca^2+^-free states (**Supplementary Figure 3**). However, unlike R-GECO1 (t_1/2_ = 0.56 s for Ca^2+^-free state)^28^), the photoactivation rise and decay of FR-GECOs happens within 10s of milliseconds. This fast rise and decay suggests that the photoactivation behaviour of FR-GECOs, while not desirable, could be easily distinguished from action potential-evoked GECI Ca^2+^ fluorescence changes, which typically happens on the timescale of seconds due to the response kinetics of GECIs.

### Performance in cultured neurons

The *in vitro* characterization of FR-GECO1a and FR-GECO1c revealed that, fluorescence colour aside, these two new GECIs had properties, such as large fluorescence changes and high Ca^2+^ affinity, comparable to state-of-the-art GECIs. Of the two indicators, we expected that FR-GECO1a might be more suitable for labelling fine subcellular neuronal processes, such as axons and dendrites, due to its brighter Ca^2+^-free state as shown by the characterization *in vitro*. On the other hand, we expected that FR-GECO1c might provide better performance for the detection of single spikes in neurons in terms of fluorescence change because of its larger Ca^2+^-dependent Δ*F*/*F*_0_ *in vitro* relative to FR-GECO1a. To evaluate the performance of FR-GECO1a and FR-GECO1c for imaging of neuronal activity, we expressed each GECI in dissociated rat hippocampal neurons. We observed that the fluorescence in neurons expressing either FR-GECO1 variant was evenly distributed throughout the cytosol (**Figure 2A, B**). Of the two indicators, FR-GECO1a did, indeed, have a brighter basal fluorescence, providing easier identification of transfected cells as well as facilitating visualization of the fine morphology of neural cells.

**Figure 2.**
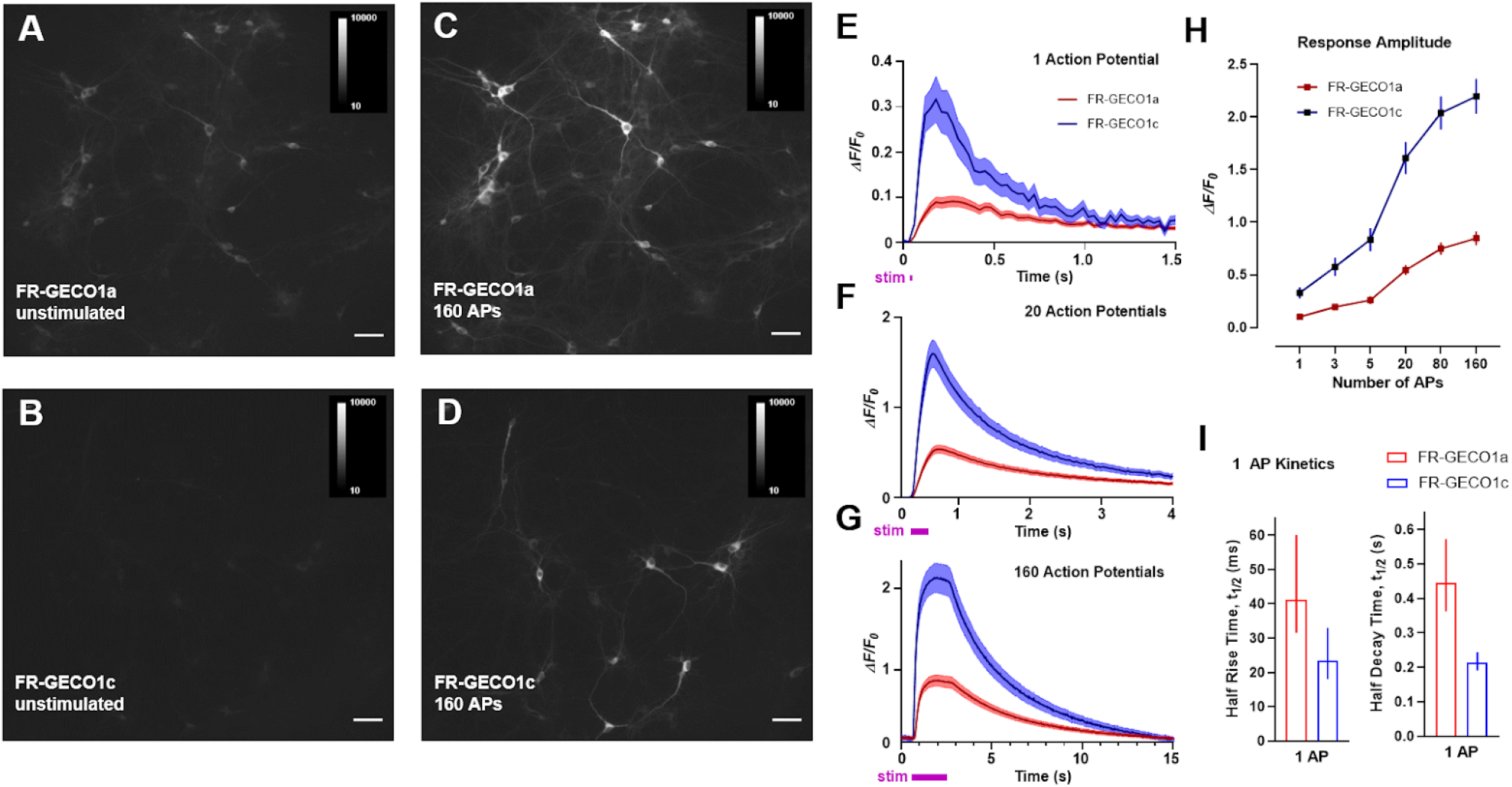
Performance of FR-GECO1 in cultured neurons. Representative images of neurons with FR-GECO1a at resting state (**A**), FR-GECO1c at resting state (**B**), FR-GECO1a at stimulated state (**C**), and FR-GECO1c at stimulated state (**D**). Averaged traces of FR-GECO1a (blue) and FR-GECO1c (red) recorded from neurons for 1 action potential (**E**), 20 action potentials (**F**) and 160 action potentials (**G**). Traces were normalized to baseline. Shaded areas represent S.E.M. in E, F and G. **H**. Averaged responses of FR-GECO1a (blue) and FR-GECO1c (red) as a function of the number of action potentials. **I**. Rise and decay time of FR-GECO1a and FR-GECO1c fluorescence changes induced by a single AP. Error bars represent S.E.M. in H and I. (n = 68 neurons for FR-GECO1a and n = 81 neurons for FR-GECO1c).

To evaluate indicator function, we acquired fluorescence images while delivering electric field stimuli to evoke action potentials (APs) (in trains of 1, 2, 3, 5, 10, 20, 40, 80, 120, 160 APs) (**Figure 2C, D**). We observed that a single stimulus elicited a 10% Δ*F*/*F*_0_ response for FR-GECO1a (n = 81 neurons) and a 33% Δ*F*/*F*_0_ for FR-GECO1c (n = 68 neurons) (**Figure 2E**). FR-GECO1c’s response to a single stimulus is ~50% higher than reported responses for jR-GECO1a and K-GECO1, suggesting that FR-GECO1c is among the most sensitive RFP-based GECIs for single AP detection^16–18^). FR-GECO1a and FR-GECO1c showed Δ*F*/*F*_0_ of 48% and 158%, respectively, upon 20 stimuli by the field electrode (**Figure 2F**). When evoked with 160 stimuli, neurons expressing FR-GECO1a exhibited a Δ*F*/*F*_0_ of 85% while FR-GECO1c had a Δ*F*/*F*_0_ of 220% (**Figure 2G, H**). In terms of single AP response kinetics, consistent with *in vitro* kinetics measurement, both indicators exhibited relatively fast rise (half rise time, FR-GECO1a: 41 ms; FR-GECO1c: 23 ms) and decay (half decay time, FR-GECO1a: 0.44 s; FR-GECO1c: 0.21 s) times, with FR-GECO1c having ~2-fold faster rise and decay kinetics than FR-GECO1a (**Figure 2I**). Overall, these results suggest that both FR-GECO1a and FR-GECO1c enable sensitive detection of neuronal activities. FR-GECO1c, in particular, offers exceptional single AP detection sensitivity, fast response times, and large fluorescence changes, compared to other state-of-art red GECIs^16–18^).

## Discussion

GFP- and RFP-based GECIs have been extensively engineered and are widely used for imaging of cell signalling and neural activity in mammalian tissue *in vivo*. However, there remains a need for GECIs with more red-shifted wavelengths, such as indicators with wavelengths in the far-red or near-infrared range. Indicators with longer excitation and emission wavelengths enable deeper imaging, which may be useful for imaging subcortical structures. Red-shifted indicators are also generally preferable as they provide better spectral separation from green indicators and commonly used optogenetic tools. They also show less phototoxicity and autofluorescence. Here, we have reported the engineering of FR-GECO1a/c, the first far-red fluorescent GECIs that are analogous to the autofluorescent protein-based GCaMP series of Ca^2+^ indicators. Our initial effort to engineer a far-red Ca^2+^ indicator using other fluorescent protein scaffolds were unsuccessful, likely due to these scaffolds’ oligomerization tendencies. On the other hand, mKelly2 is reported to be a true monomeric far-red fluorescent protein, a property which likely enabled the engineering of the first far-red fluorescence fully genetically-encoded indicator. Moreover, we expect that the cpmKelly scaffold of FR-GECO1 can serve as a template for engineering new FP-based far-red indicators for other ligands.

Both FR-GECO1 variants were also bright relative to other red GECIs under 2P conditions. Their optimal 2P Ca^2+^-dependent Δ*F*/*F*_0_ occurs between 1050 and 1200 nm, overlapping with the range of common tunable 2P lasers such as titanium-sapphire lasers ranging from 650 to 1100 nm. When excited at 1200 nm, FR-GECO1a and FR-GECO1c, in their Ca^2+^-saturated states, have very high 2P brightness. At this wavelength, their brightness is comparable to a handful of bright far-red FPs, such as mKates, Katushkhas, and Neptunes. Both of these sensors are much brighter upon 2P excitation at 1200 nm than any of the red GECIs characterized so far^24^) (**Supplementary Table 3**). Presumably, 2P microscopy at 1200 nm excitation will provide deeper focusing ability compared to 1100 nm excitation because of less scattering. Although water absorbs stronger at 1200 nm, the depths of imaging are not too large (typically up to 200 – 300 μm), and this would not cause any attenuation of the laser beam. However, heating due to the laser energy absorption by water could be stronger. Improved 2P brightness of the FR-GECOs should help reduce the required laser power for excitation, and consequently, tissue heating^29,30^.

NIR-GECO1 has excitation and emission peaks at 678 and 704 nm, respectively, with a 10-fold inverted response in fluorescence intensity upon Ca^2+^ addition with a *K*_d_ of 215 nM. Although not as red-shifted as NIR-GECO1, FR-GECO1 variants have higher molecular brightness, and have a positive response to Ca^2+^. Indicators with a positive response to their ligands are generally favoured over indicators with an inverse response due to a better signal-to-noise ratio owing to lower background fluorescence. In addition, both FR-GECO1a and FR-GECO1c offer better sensitivity (Δ*F*/*F*_0_ = 10% and 33% for 1AP and 85% and 220% for 160 APs for FR-GECO1a and FR-GECO1c, respectively) and faster kinetics (half rise time of 41 ms and 23 ms and half decay time of 440 and ~210 ms for 1 AP for FR-GECO1a and FR-GECO1c, respectively) than NIR-GECO1 (−Δ*F*/*F*_0_ = 2.3% for 1 AP and 48% for 160 APs with 0.94 s rise time and 2.8 s decay time for 1 AP and 1.3 s rise time and 6.5 s decay time for 160 APs) in neuronal activity detection.

In addition to the unique spectral profile, the FR-GECO1 series has other advantages relative to other green and red GECIs. FR-GECO1c exhibits a larger Ca^2+^-dependent Δ*F*/*F*_0_ *in vitro* than most other red GECIs, such as R-GECO1, K-GECO1, and XCaMP-R^17,18,21^. The use of ckkap peptide provides FR-GECO1a and FR-GECO1c with strong affinities for Ca^2+^ (*K*_d_ <100 nM), as well as large fluorescence changes and fast response times in single AP detection, similar to K-GECO1 and the XCaMP series GECI, when compared against other RS20-based indicators such as R-GECO1, and the GCaMP and RCaMP1 series^17,18,21,28,31^. In particular, FR-GECO1c offers better Δ*F*/*F*_0_ in single action potential detection in cultured neurons (Δ*F*/*F*_0_ = 33% for 1AP) than FR-GECO1a and all the other aforementioned red GECIs. This excellent sensitivity of FR-GECO1c is likely due to its lower Ca^2+^-free state fluorescence, larger Ca^2+^-dependent Δ*F*/*F*_0_, and optimal affinity towards Ca^2+^.

As first generation sensors, FR-GECO1a and FR-GECO1c both still have room for further improvements. A general trend among FPs is that hypsochromic shifts in peak wavelengths are accompanied by reductions in quantum yield, thus limiting the brightness of available far-red and near-infrared FPs. Future versions of FR-GECO should be optimized for improved brightness to further facilitate imaging *in vivo*, and even faster kinetics. Given that our pH titrations revealed that both FR-GECO1 variants have a significantly more basic p*K*_a_ relative to mKelly2 and do not exhibit their maximum brightness under physiological conditions, engineering variants with lower p*K*_a_ values may also lead to improved brightness for imaging in tissue. Additionally, the red-shifted spectrum of FR-GECO1a/c makes them excitable with wavelengths beyond 600 nm. Although not optimal, FR-GECO1a/c are the only non-biliverdin-binding genetically encoded Ca^2+^ indicator that can be excited by the widely used red helium-neon lasers (~633 nm, ~12.5% of max excitation). We thus believe further red-shifted FR-GECO1a/c with better compatibility with helium-neon lasers will also be of interest to researchers.

In summary, we have developed far-red genetically-encoded indicator Ca^2+^ indicators, the FR-GECO series, expanding the spectral palette of available GECIs that utilize non-biliverdin-binding fluorescent proteins. At its first iteration, FR-GECO1a/c showed good brightness, strong affinity for Ca^2+^, and good sensitivity for neuronal activity detection. In the future, we aim to improve its brightness and demonstrate its application *in vivo*. Furthermore, we expect that the cpmKelly2 scaffold can be used to further expand the color palette of single FP-based genetically encodable indicators for a wide range of analytes (i.e., other than Ca^2+^)^32,33^. Thus, we believe that FR-GECO1 is a valuable new addition to the GECI toolkit.

## Methods

### General

Synthetic DNA oligonucleotides and double-stranded fragments (gBlocks) were sourced from Integrated DNA Technologies. CloneAmp HiFi PCR Premix and Q5 polymerase were used for high fidelity polymerase chain reactions (PCRs), and were purchased from Takara Bio USA, Inc. and New England BioLabs, respectively. Taq polymerase was purchased from Thermo Fisher Scientific. PCR products were purified by preparative agarose gel electrophoresis and extracted with GeneJET Gel Extraction Kit purchased from Thermo Fisher Scientific. Restriction enzymes and T4 DNA ligase were purchased from Thermo Fisher Scientific and used according to the manufacturer’s recommended protocols. Digested PCR products were extracted with the GeneJET Gel Extraction Kit before ligation. In-Fusion HD was purchased from Takara Bio USA, Inc. and used for assembly reactions. Plasmid DNA extractions were performed with GeneJET Plasmid Miniprep Kits (Thermo Fisher Scientific). Sanger sequencing reactions were performed by the Molecular Biology Services Unit at the University of Alberta. Fluorescence measurements were performed on a Safire^2^) plate reader (Tecan), while absorbance measurements were collected on a DU-800 UV-Visible Spectrophotometer (Beckman).

### Protein Engineering

#### Starting template construction

Double-stranded DNA encoding for K-GECO1 (Ref. 17), with its circularly permuted (cp) fluorescent protein replaced with cpmKelly2 (Ref. 12), was commercially synthesized with overlapping sequences with pBAD/HisB (Thermo Fisher Scientific) in the 5’ and 3’ directions to facilitate insertion between the XhoI and HindIII sites. This fragment was then assembled into pBAD/HisB vector digested with XhoI and HindIII, transformed into electrocompetent DH10B *E. coli* (Invitrogen), and incubated overnight on Lennox Broth (LB) agar plates supplemented with 400 μg/mL ampicillin (Thermo Fisher Scientific) and 0.02% (w/v) L-arabinose (Alfa Aesar) at 37°C. Plasmids were isolated from single colonies grown in 4 mL liquid LB cultures with 100 μg/mL ampicillin, and sequenced by Sanger sequencing.

#### Plasmids for mammalian expression

Genes encoding FR-GECO1 and FR-GECO1c were amplified by high-fidelity PCR using primers that add vector homologous sequences to the 5’ and 3’ end and assembled into linearized pcDNA3.1/Puro-CAG-ASAP1 to replace the gene encoding for ASAP1 (a gift from Michael Lin, Addgene plasmid #52519)^34^).

#### Library construction

To generate random mutagenesis libraries, mutations were introduced into the construct by error-prone PCR amplification with Taq polymerase in the presence of MnCl_2_ (varies; up to 0.15 mM) and a 1:5 ratio of dATP and dGTP to dCTP and dTTP. The purified PCR product was then assembled with a *Xho*I and *Hind*III digested pBAD/HisB vector. Site-directed mutagenesis libraries were generated by PCR amplification with high fidelity polymerases and primers carrying the codons for the desired mutations followed by DpnI digestion. The NNK codon was used for complete randomization. For mutations at just one position, amplification was performed with the CloneAmp HiFi PCR Premix, while libraries with mutations at multiple locations were generated by QuikChange Lightning Multi Site-Directed Mutagenesis Kit (Agilent Technologies).

#### Library screening

Libraries were transformed into DH10*B E. coli* and cultured on LB agar plates supplemented with ampicillin and L-arabinose, as described earlier. The *E. coli* colonies expressing the library were then screened on the plate on the basis of fluorescence intensity using a custom imaging system described previously^35^). Colonies displaying the highest fluorescence intensity were then cultured in 4 mL liquid LB with ampicillin and arabinose overnight at 37°C. Proteins were then extracted from cells using B-PER (Thermo Fisher Scientific) and subjected to a secondary screen by measuring their fluorescence intensities in Ca^2+^-free (30 mM 3-(N-morpholino)propanesulfonic acid (MOPS), 100 mM KCl, 10 mM ethylene glycol-bis(β-aminoethyl ether)-*N,N,N’,N’*-tetraacetic acid (EGTA), pH 7.2) and Ca^2+^ buffers (30 mM MOPS, 100 mM KCl, 10 mM Ca-EGTA, pH 7.2) to determine their Ca^2+^-dependent Δ*F*/*F*_0_. Plasmids for variants showing the largest Ca^2+^-dependent Δ*F*/*F*_0_ and highest fluorescence intensities were extracted, sequenced, and used as the template(s) for the next round of evolution.

### *In vitro* characterization

#### Protein expression and purification

The pBAD/HisB plasmid carrying the gene encoding for the protein of interest was transformed into electrocompetent DH10B *E. coli* and grown on solid media. Single colonies from the transformed bacteria were then used to inoculate 4 mL of a starter culture supplemented with ampicillin and L-arabinose that was then shaken at 225 rpm in 37°C overnight. The starter culture was added to 500 mL of LB with 100 μg/mL ampicillin and the culture was shaken at 37°C. After 4 h, 0.02% L-arabinose is added to induce expression, and the culture is shaken for another 4 h at 37°C before harvesting the bacteria by centrifugation. Bacteria were then resuspended in 1× TBS (50 mM Tris-HCl, 150 mM NaCl, pH 7.5), and lysed by sonication. The lysate was clarified by centrifugation and the cleared lysate was incubated with Ni-NTA resin for at least 1h. Resin bound to protein was washed with TBS wash buffer (1× TBS with 20 mM imidazole, pH 8.0) followed by elution with TBS elution buffer (1× TBS with 250 mM imidazole, pH 7.8). The purified protein was then left at room temperature overnight to facilitate protein folding and chromophore maturation before concentrating and buffer exchanging into 1× TBS using 10 kDa centrifugal filter units (Millipore). Unless noted otherwise, all steps were carried out at 4°C or on ice.

#### Brightness, Affinity, and pH-sensitivity measurement

Effective ECs of FR-GECO variants were measured using the alkali denaturation method [4]. Briefly, purified protein for each FR-GECO variant was diluted in NaOH, Ca^2+^-free (30 mM MOPS, 100 mM KCl, 10 mM EGTA, pH 7.2), and Ca^2+^ buffer (30 mM MOPS, 100 mM KCl, 10 mM Ca-EGTA, pH 7.2), and the absorption spectrum for each sample was collected. Because the same amount of protein was used for each set of measurements for each protein, the concentration of protein present in each sample can be calculated according to Beer’s Law and the previously reported EC of 44,000 for the absorbance peak around ~ 450 nm, which corresponds to the denatured tyrosine-based chromophore^23,36^. Further application of Beer’s Law with the calculated concentration for each variant yields the EC in the Ca^2+^-bound and Ca^2+^-free states. Measurements were performed in triplicate and the results were averaged. Determination of QY was performed according to previously established protocols using mKelly2 as the standard^12,21,35^. For each protein, a series of diluted samples were prepared in Ca^2+^-free and Ca^2+^ buffer from the samples used for extinction coefficient measurements such that the peak absorbance was equal to, or less than, 0.05. For each dilution series, the emission spectra were collected from 590 to 750 nm with an excitation wavelength of 570 nm. The total fluorescence intensities for each dilution were integrated, plotted against the absorbance, and the slope of each line (*m)* was calculated. Quantum yields were then calculated using the published quantum yield for mKelly2 (0.18; Ref. 12) and according to the equation:

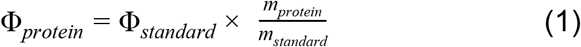

Buffers for the *K*_d_ determination, with free [Ca^2+^] ranging from zero to 39 μM, were prepared by mixing the appropriate volumes of Ca^2+^-free and Ca^2+^ buffers, as described previously^37^). Purified FR-GECO variants were diluted in these buffers and the fluorescence intensities of the protein in each solution were measured in triplicate. The readings were plotted against the free Ca^2+^ concentration on a logarithmic scale, and the data fitted to the Hill equation for the *K*_d_ and apparent Hill coefficient. Aliquots of a buffer containing 30 mM trisodium citrate and 30 mM borax (sodium borate) were adjusted using HCl or NaOH so that their pH ranged from 3 to 11. Concentrated protein, as well as 1 μL of either 200 mM EGTA or CaCl_2_ for the Ca^2+^-free and Ca^2+^ conditions, respectively, were then diluted in these buffers for a total volume of 100 μL. The fluorescence intensities were measured in triplicate, and the readings were plotted against pH. Due to the ionization events at pH ~ 5 to 6, only the data for pH 6.5 to 11 were fitted to the Hill equation for the p*K*_a_.

#### *In vitro* kinetics measurement

Rapid kinetic measurements of FR-GECO1a and FR-GECO1c in purified protein were made using an Applied Photophysics SX-20 Stopped-Flow Reaction Analyzer using fluorescence detection. The dead time of the instrument was 1.1 ms. The excitation wavelength used was 625 nm with 2 nm bandwidth and emitted light was collected at 690 nm through a 10 mm path at room temperature. For *k*_off_ measurements, 2 μM of each FR-GECO1a and FR-GECO1c proteins in 1 mM Ca^2+^ (30 mM MOPS, 100 mM KCl, pH 7.2) were rapidly mixed at 1:1 ratio with 100 mM of EGTA (30 mM MOPS, 100 mM KCl, pH 7.2) at room temperature. Measurements were taken for five replicates. *k*_off_ values were determined by fitting a single exponential dissociation curve to the signal decay with units of s^−1^,using Prism GraphPad. For *k*_on_, both variants, buffered in 30 mM MOPS and 100 mM KCl (pH = 7.2), were rapidly mixed at 1:1 ratio with varying concentrations of Ca^2+^ produced by reciprocal dilutions of 10 mM EGTA and 10 mM CaEGTA by using the Calcium Calibration Buffer Kit (ThermoFisher Scientific). The slope of *k*_obs_ was used to determine *k*_on_ rate in the units of s^−1^ M^−1^. Two replicates were used for each Ca^2+^ concentration.

#### Photoactivation measurement

Microdroplets of FR-GECO1a and FR-GECO1c in purified protein in either Ca^2+^ or Ca^2+^-free buffer (pH 7.2) were prepared using Octanol. The prepared sample was sandwiched between pre-silanized coverglass and coverslip. Using a widefield microscope each droplet was continuously illuminated with a 561 nm laser for 120 sec. 488 nm laser was used for photoactivation while the 561 nm laser was on. The 488 nm laser was on for 50 ms with the shutters switching on at various frequencies (0.5, 1, or 2 Hz).

#### 2P characterization

Experimental setup for 2P spectral measurements includes a tunable femtosecond laser InSight DeepSee Dual (Spectra Physics) coupled with a photon counting spectrofluorometer PC1 (ISS)^24,38^. The 2P fluorescence excitation (2PE) spectra were measured by automatically stepping laser wavelength and recording total fluorescence intensity at each step. A combination of filters, including FF01-770/SP, FF01-680/SP (both Semrock), and FGL630 long-pass (Thorlabs) was used in the left emission channel of PC1 spectrofluorometer to eliminate scattered laser light. To scale the 2P excitation spectra, we use a parameter called 2P molecular brightness *F*_2_(*λ*). This parameter depends on fractional contributions to 2P brightness of the anionic and neutral forms at excitation wavelength *λ* and can be presented as follows:

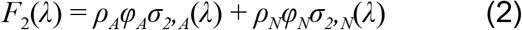

where *φ* is the fluorescence quantum yield of form *A* or *N*, and *σ*_*2*_ is the 2P absorption cross section. Since we collect fluorescence spectrum through a FGL630 long-pass filter, technically *φ*_*N*_ only comprises a part of fluorescence originating from anionic deprotonated chromophore that appears after the excited-state proton transfer from initially excited neutral form. To obtain the 2P excitation spectrum in units of molecular brightness, we independently measured *ρ*_*A*_, *φ*_*A*_, and *σ*_*2,A*_(λ) of the anionic form for both FR-GECO1a/c in Ca^2+^-free and Ca ^2+^-saturated states and normalized the unscaled 2P spectrum to the product *ρ*_*A*_ *φ*_*A*_ *σ*_*2,A*_(*λ*).

The cross section *σ*_*2,A*_(*λ*) was measured at the wavelength where the contribution of the neutral form is negligible (*λ* = 1064 nm). This measurement was performed using Rhodamine B in alkaline ethanol as a reference standard. Its cross section was measured relatively to Rhodamine 6G in ethanol (with the average <*σ*_*2*,RhB_/*σ*_*2*,Rh6G_> = 1.072 ± 0.017). Using this number together with the Rhodamine 6G cross section in ethanol (*σ*_*2*,Rh6G_) obtained after averaging of seven independent measurements^38^): <*σ*_*2*,Rh6G_> = 12.4 ± 2.6 GM, cross section of Rhodamine B in ethanol at 1064 nm was calculated *σ*_*2*,RhB_ = 13.3 ± 2.8 GM and used as a reference. In a modified version of previously reported method^38^), a total (without monochromator) 2P excited (at 1064 nm) fluorescence signals *I* as a function of laser power *P* were collected for both the sample and reference solutions (solutions were held in 3×3 mm cuvettes (Starna) with maximum optical density less than 0.1). The fluorescence was collected at 90° to excitation laser beam through FF01-770/SP, FGL630 Longpass, and long wave pass 561 (Edge Basic™, Semrock) filters, using the left emission channel of a PC1 spectrofluorometer working in a photon counting mode. The power dependences of fluorescence were fit to a quadratic function *I* = *aP*^2^), from which the coefficients *a*_S_ and *a*_R_ were obtained for the sample (index S) and reference (index R) solutions, respectively. Second, the 1P excited fluorescence signals were measured for the same samples and in the same registration conditions. In this case, a strongly attenuated radiation of a 561 nm line of a Sapphire 561-50 CW CDRH (Coherent) laser was used for excitation. The fluorescence power dependences for the sample and reference were measured and fit to a linear function: *I* = *bP*, from which the coefficients *b*_*S*_ and *b*_*R*_ were obtained. The 2P absorption cross section was then calculated as follows:

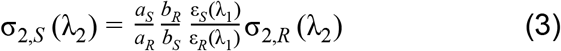

Here, λ_1_ is the wavelength used for 1P excitation (561 nm), λ_2_ is the wavelength used for 2P excitation (1064 nm), *ε*_*R,S*_(λ_1_) are the corresponding extinction coefficients, measured at λ_1_. (In contrast to the one-photon characterization section, we define extinction coefficient (ε_S_) here to be of a particular, anionic, form. It relates to the previously defined extinction coefficient, e.g. shown in Table 1, as follows: ε_S_ = ε/ρ_A_) This approach allows us to automatically correct for the laser beam properties (pulse duration and spatial intensity distribution), fluorescence collection efficiencies for 1P and 2P modes, PMT spectral sensitivity, differences in QYs and concentrations between S and R solutions. Molecular brightness of the anionic form was then calculated as a product *F*_2_ = *ρ*_*A*_*φ*_*A*_*σ*_*2,A*_(*λ_m_*) with the 2P cross section taken at spectral maximum, *λ*_m_, for both states of the sensor. The QYs, ECs, and fractional concentrations were all measured independently. Finally, the 2P excitation spectra were scaled to the calculated *F*_2_(1064 nm) values.

### Imaging in cultured neuron

Primary rat hippocampal neurons were prepared as described previously^39^) Cultured neurons were transfected with pCAG-FR-GECO1a and pCAG-FR-GECO1c plasmids using electroporation (Lonza Nucleofector) according to manufacturer’s instructions. Prior to imaging session, transfected neurons were first washed twice with imaging buffer (145 mM NaCl, 2.5 mM KCl, 10 mM glucose, 10 mM HEPES, pH 7.4, 2 mM CaCl_2_ and 1 mM MgCl_2_), and treated with synaptic blockers (10 μM CNQX, 10 μM CPP, 10 μM GABAZINE, and 1 mM MCPG). Neurons were imaged using an inverted Nikon Eclipse Ti2 microscope equipped with a SPECTRA X light engine (Lumencor), a 20× objective (NA = 0.75, Nikon), and a sCMOS camera (Hamamatsu ORCA-Flash 4.0). Neurons were stimulated by field stimulation with a custom-built platinum wire electrode with a stimulus isolator (A385, World Precision Instruments). Trains of 1, 2, 3, 5, 10, 20, 40, 80, 120 and 160 APs were stimulated and fluorescence images were acquired throughout the stimulation session. Acquired images were processed with ImageJ.

## Supporting information

Supplementary Video 1. Performance of FR-GECO1a in cultured neurons.

Supplementary Video 2. Performance of FR-GECO1c in cultured neurons.

## Author contributions

YS, REC conceived the project. YS, RD, JW performed protein engineering. RD, YS, MD, RSM, AA, RP performed *in vitro* protein characterization. YS, ASA, YZ performed cultured cell imaging. YS, KP, ERS, TEH, REC supervised the research. YS, RD, MD, REC wrote the manuscript. All authors read and approved the final manuscript.

## Acknowledgments

We thank the University of Alberta Molecular Biology Services Unit for technical support. We thank Dr. Timothy Wannier for sharing mKelly sequences. We thank Dr. Fern Sha, Rosario Valenti, and the Janelia Research Campus cell culture facility for technical assistance. Ronak Patel is a member of the Janelia Research Campus Tool Translation Team (T3); we thank the T3 for providing equipment access and technical support. This work was supported by grants from the Natural Sciences and Engineering Research Council of Canada (NSERC; RGPIN 2018 04364), the Canadian Institutes of Health Research (CIHR; FS 154310), and Janelia Visiting Scientist Program (Janelia Research Campus, HHMI).

## Material and data availability

The data supporting this research are available upon request. Plasmid constructs encoding FR-GECO1a and FR-GECO1c are available through Addgene.

## Supplementary information

**Supplementary Figure 1.**
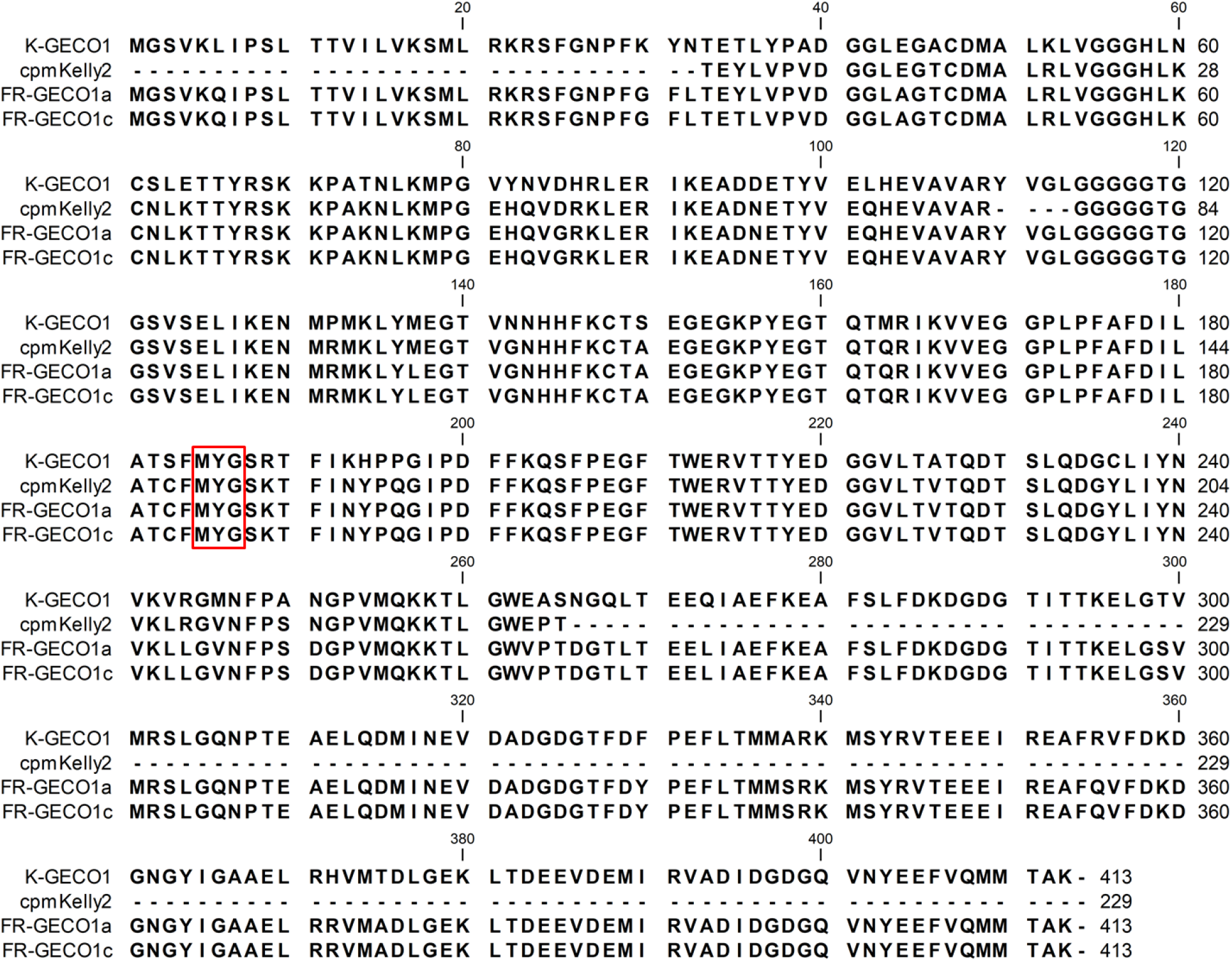
Sequence alignment of K-GECO1, mKelly2, FR-GECO1a, and FR-GECO1c. Chromophore-forming residues are indicated by a red box.

**Supplementary Figure 2.**
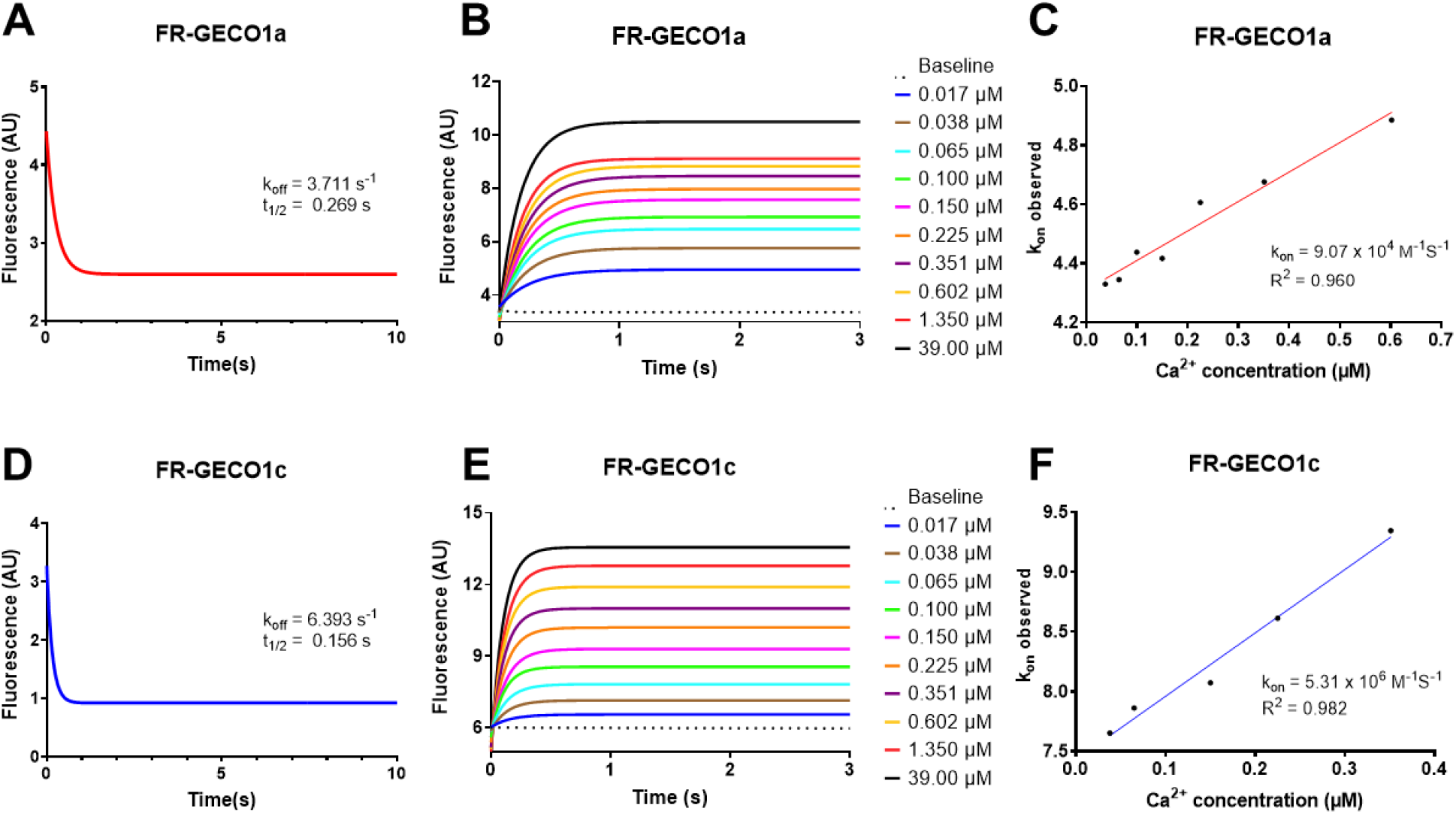
Kinetics of FR-GECO1a and FR-GECO1c purified proteins. Ca^2+^ dissociation kinetics of FR-GECO1a (**A**) and FR-GECO1c (**D**). Ca^2+^ association kinetics of FR-GECO1a (**B**) and FR-GECO1c (**E**). Observed rate constants plotted as a function of Ca^2+^ concentration for FR-GECO1a (**C**) and FR-GECO1c (**F**).

**Supplementary Figure 3.**
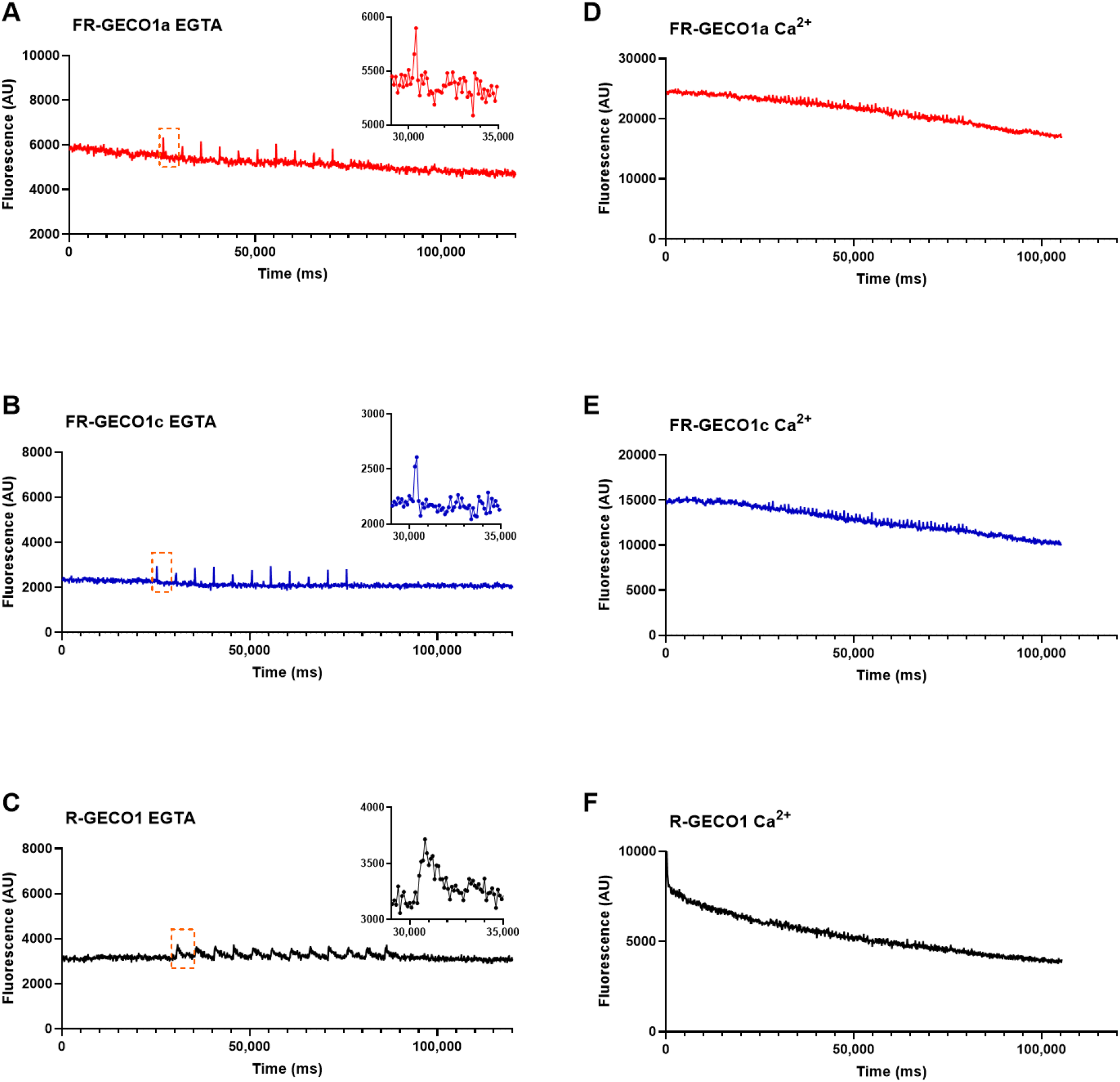
*In vitro* photoactivation of FR-GECO1a, FR-GECO1c, and R-GECO1 purified proteins. Representative fluorescence responses of FR-GECO1a (**A**), FR-GECO1c (**B**), and R-GECO1 (**C**) to 50 ms of pulse (0.5 Hz) illumination with a 488 nm (0.71 W/cm^2^)) laser and constant illumination with a 561 nm (3.70 W/cm^2^)) laser in the absence of Ca^2+^ (MOPS-EGTA buffer). Insets: zoom-in view of single photoactivation event traces. Representative FR-GECO1a (**D**), FR-GECO1c (**E**), and R-GECO1 (**F**) fluorescence responses to 50 ms of pulse (1 Hz) illumination with a 488 nm (0.71 W/cm^2^) laser and constant illumination with a 561 nm (3.70 W/cm^2^) laser in the presence of Ca^2+^ (MOPS-CaEGTA buffer).

**Supplementary Table 1.**
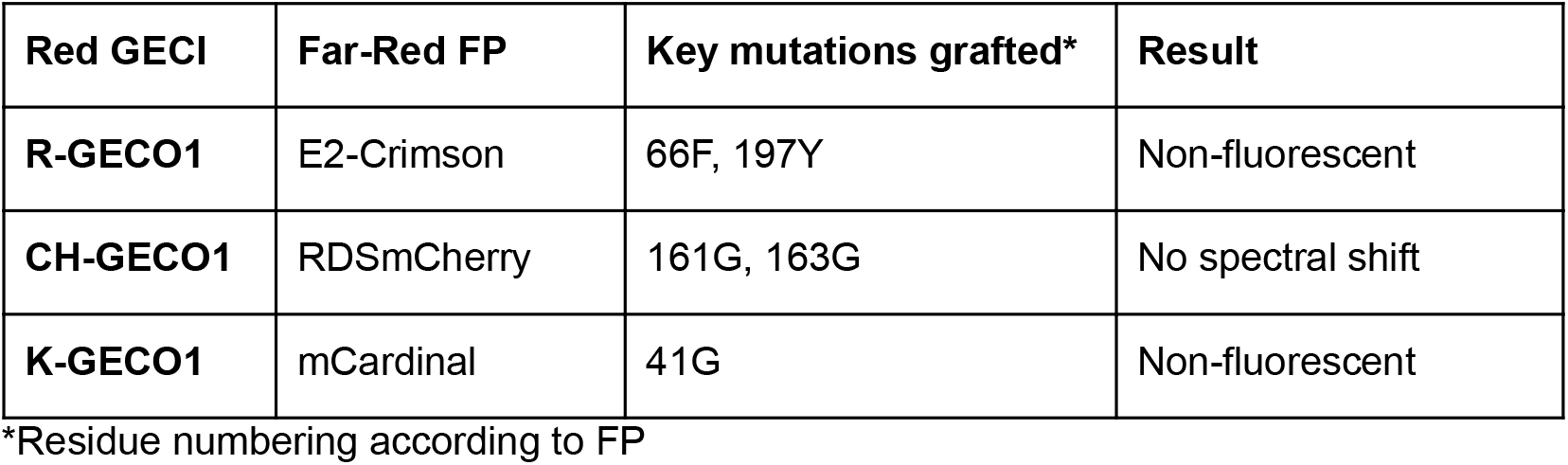
Key mutations of far-red FPs grafted onto existing red GECIs

| Red GECI | Far-Red FP | Key mutations grafted* | Result |
| --- | --- | --- | --- |
| **R-GECO1** | E2-Crimson | 66F, 197Y | Non-fluorescent |
| **CH-GECO1** | RDSmCherry | 161G, 163G | No spectral shift |
| **K-GECO1** | mCardinal | 41G | Non-fluorescent |

\*Residue numbering according to FP

**Supplementary Table 2.** Mutations accumulated during engineering of FR-GECOs.

| Name | Mutations (ckkap Linker cpmKelly2 CaM) |
| --- | --- |
| 0.1 | Template |
| 0.2 | 0.1 + K30G Y31F N32L |
| 0.3 | 0.2 + (M1V) D85G N251D R355Q H372R |
| 0.4 | 0.3 + E263V |
| 0.5 | 0.4 + R244L |
| 0.6 | 0.5 + Q268R Q273L |
| 0.7 | 0.6 + L6Q M137L N266D |
| 0.8 | 0.7 + (V1M) T299S A338S T375A |
| 0.9 | 0.8+ E44A F330Y |
| 1a | 0.9 + R268T |
| 1c | 1a + Y35T |

**Supplementary Table 3.** *In vitro* 2P comparison of FR-GECOs with other red GECIs.

| Red GECI | 2P brightness of<br>Ca<sup>2+</sup>-saturated form<br>of the anionic<br>chromophore ($\varphi \times \sigma_2$)<br>GM (at $\lambda_{2P,max}$) | 2P brightness of<br>Ca<sup>2+</sup>-saturated form<br>of the anionic<br>chromophore ($\varphi \times \sigma_2$)<br>GM (1200 nm) | $F_2(\lambda_{2P,max})$<br>($\varphi \times \sigma_2 \times \rho$)<br>GM | $F_2(1200\text{ nm})$<br>($\varphi \times \sigma_2 \times \rho$)<br>GM |
| --- | --- | --- | --- | --- |
| FR-GECO1a | 20 (1112 nm) | 15 | 9.2 | 6.9 |
| FR-GECO1c | 15 (1112 nm) | 11 | 7.4 | 5.5 |
| R-GECO1.2 | 9.6 (1044 nm) | 0.13 | 8 | 0.10 |
| CAR-GECO1 | 9.4 (1060 nm) | 0.6 | 8 | 0.54 |
| K-GECO1 | 11 (1068 nm) | 1.1 | 8 | 0.83 |
| jRCaMP1a | 12.4 (1076 nm) | 2.7 | 10 | 2.2 |

\* Data for R-GECO1.2, CAR-GECO1, K-GECO1, and jRCaMP1a from Molina et al.<sup>24</sup>

**Supplementary Video 1.** Performance of FR-GECO1a in cultured neurons.

**Supplementary Video 2.** Performance of FR-GECO1c in cultured neurons.

